# Aggregation of an FG nucleoporin under crowded conditions

**DOI:** 10.1101/2024.04.15.589310

**Authors:** Laura Maguire, Sophie Reskin, Kathryn Wall, Elena Arroyo, Paul Marchando, A. M. Whited, Annette Erbse, Steven T. Whitten, Loren E. Hough

**Affiliations:** Department of Physics, University of Colorado Boulder, Boulder, Colorado, United States of America; BioFrontiers Institute, University of Colorado Boulder, Boulder, Colorado, United States of America; Department of Biochemistry, University of Colorado Boulder, Boulder, Colorado, United States of America; Department of Chemistry and Biochemistry, Texas State University, San Marcos, Texas 78666, United States

## Abstract

Macromolecular crowding can affect the aggregation behavior of intrinsically disordered proteins in unexpected ways. We studied the aggregation of a peptide derived from the disordered FG nucleoporins which line the nuclear pore complex. We measured its aggregation kinetics in the presence of both inert and non-specifically interacting crowding agents. Using fluorescence emission and NMR spectroscopy, we probed differences in the local chemical microenvironment of the peptide’s residues. Our results indicate differences in aggregation kinetics and residue microenvironment depending on the identity of the crowder, including differences between crowding with PEG and PVP, two polymers which are often used interchangeably as inert crowding agents.

## Introduction

Cells are highly crowded environments, with up to 30% of their volume occupied by macromolecules [1], and this crowded environment strongly affects protein behavior. In particular, crowding often alters the aggregation properties of intrinsically disordered proteins (IDPs). Furthermore, crowding effects can differ depending on whether the crowders are inert, interacting, compact, or extended. As a general rule, crowding seems to increase the rate of aggregation of disordered proteins, but this is by no means universal [1–6]. The presence of small crowders tends to encourage proteins to take on compact conformations due to the excluded volume effect, which in the case of IDPs often leads to aggregation. However, the viscosity changes due to crowding may inhibit the formation of aggregates [7, 8], and specific chemical interactions modulate protein stability [9–11]. Crowding with proteins capable of nonspecific interactions has been shown to inhibit the partitioning of IDPs into phase-separated liquid droplets [12]. Other important but not well-understood factors include crowder structure and interactions, agitation or its absence, and the presence of an air-water interface [1, 3, 10]. As a result of these, and other effects, crowding agents differ in their effect on protein dynamics and function [**?**, 3, 6].

A system in which macromolecular crowding may be important to protein aggregation is the nuclear pore complex (NPC), which relies on highly dynamic IDPs to form a selective barrier between the cell’s nucleus and cytoplasm (Fig. 1A). Macromolecules traveling through the pore must pass a selective barrier composed of disordered FG nucleoporins (FG Nups), which consist of short hydrophobic phenylalanine-glycine (FG) motifs separated by hydrophilic linkers. Proteins above 30 kDa which do not interact with the FG motifs are largely blocked from passage. In contrast, transport factor proteins bind to the FG motifs and have high flux through the NPC [13, 14]. FG Nups exist on a continuum of cohesiveness, ranging from Nups which do not aggregation under any physiological conditions to those which do so readily [15]. Many simulations of the nuclear pore aim to understand the role of Nup cohesiveness in transport, and invoke different levels of cohesion between FG Nups [16–19].

**Figure 1.**
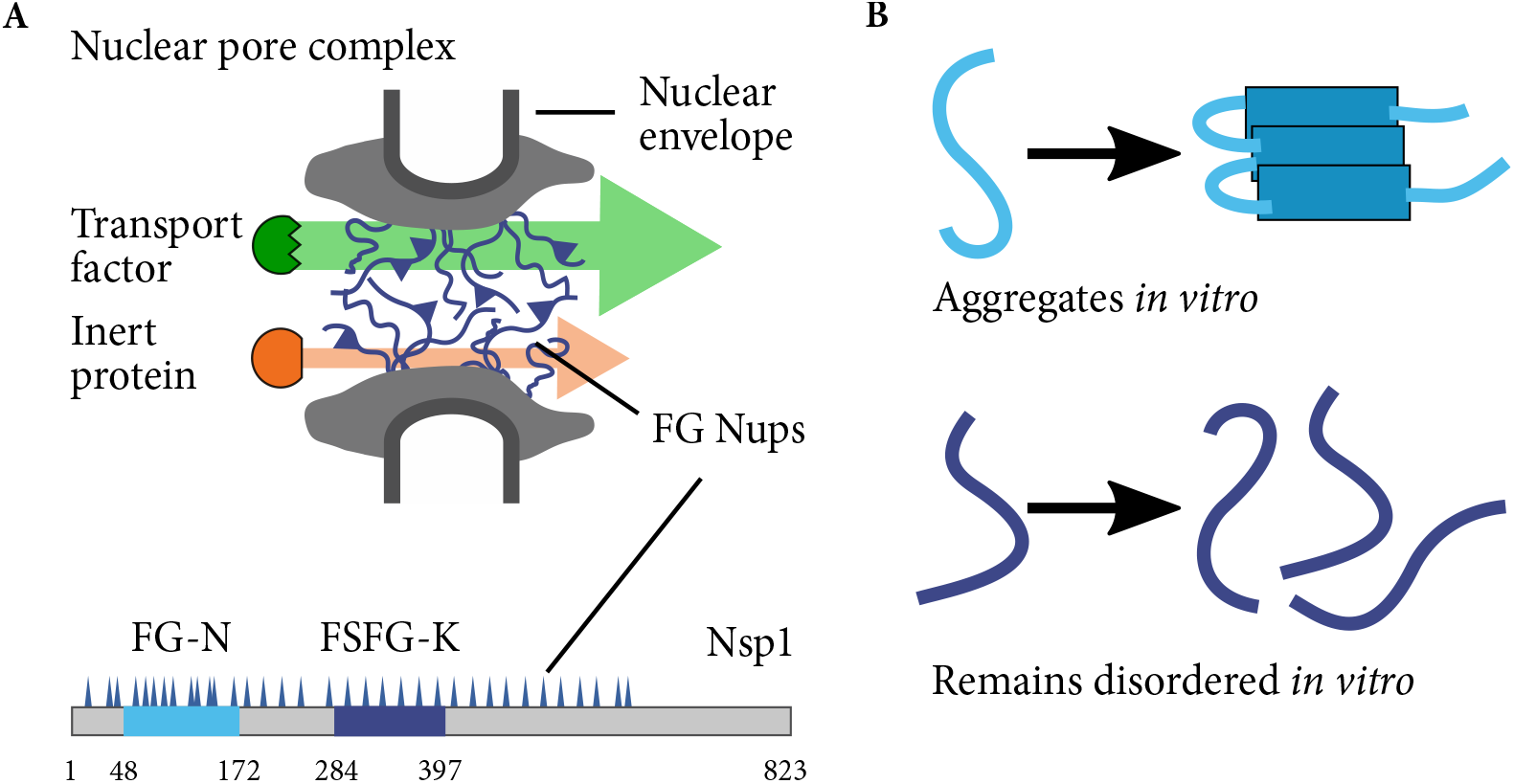
Schematics of the nuclear pore complex and FG peptides. A: The nuclear pore complex (grey) is filled with FG Nups (blue polymers) that selectively passage transport factors that bind to FG Nups (blue) while blocking non-binding proteins (red). FG-N (light blue) consists of residues 48-172 of the FG Nup Nsp1, and FSFG-K (dark blue) consists of residues 284-397. B: FG-N aggregates *in vitro* in the absence of denaturant. FSFG-K remains disordered over a wide range of buffer and pH conditions.

The conformation and aggregation states of FG Nups within the NPC are not well-known. Some Nups, such as Nsp1, aggregate in buffer to form hydrogel structures [20–22], although Nsp1 does not aggregate in the cellular environment [15]. Other Nups can form phase-separated liquid droplets [23]. Nup153 forms amyloid fibrils *in vitro* at a rate which is increased by the addition of inert crowding agents [5]. Amyloid structures are long chains of stacked *β* sheets, which are commonly formed by disordered proteins upon aggregation, and are often implicated in disease [24]. On the other hand, simulations and atomic force microscopy (AFM) data suggest that Nups in the nuclear pore remain highly dynamic [25, 26].

We tested the aggregation behavior of a well-characterized FG Nup fragment (FG-N, Fig. 1) to determine whether we could distinguish changes upon aggregation that depend on the identity of the crowding agent. The 124-amino-acid peptide FG-N is known to aggregate upon incubation in buffer, but remains disordered in cellular conditions [15]. We probed a variety of crowding agents, both inert and non-specifically interacting, using a fluorescence aggregation assay, which showed differences in the aggregation kinetics. Two inert crowders, poly(ethylene glycol) (PEG) and polyvinylpyrrolidone (PVP), were chosen for further investigation via fluorescence spectroscopy and nuclear magnetic resonance (NMR) spectroscopy. The results indicate that the presence and type of crowding agent affects not only the aggregation kinetics but also the local chemical microenvironment of the peptide’s residues.

## Materials and methods

### Protein expression and purification

We aimed to use FG Nup fragments which are similar in size, but one of which aggregates and one does not. To that end, we used the fragments FG-N and FSFG-K, which are both derived from the FG Nup Nsp1 [15]. Both FG constructs were expressed from plasmids based on pRSFDuet-1 with the second expression site removed [15]. Proteins were were expressed in BL21 DE3 Gold cells and contained a C-terminal his tag. Cultures were grown in LB to OD 0.6-0.8. Isotopically-labeled protein was expressed by pelleting cells and washing in M9 salts pH 7.4 (6.2 g/L Na_2_HPO_4_, 3.0 g/L KH_2_PO_4_, 0.5 g/L NaCl). Cells were then resuspended in M9 minimal media with ^15^N ammonium chloride and induced. Non-isotopically-labeled protein remained in LB throughout induction. All cultures were induced for 2-4 hours with 1 mM IPTG. Periplasmic matrix was removed by resuspending cell pellets in SHE buffer (20% sucrose, 50 mM HEPES, 1 mM EDTA), spinning down and resuspending in 5 mM MgSO_4_, incubating 10 minutes on ice, and spinning down and discarding supernatant [34]. Proteins were purified with a cobalt affinity column in potassium transport buffer (PTB; 150 mM KCl, 20 mM HEPES, 2 mM MgCl_2_) and eluted with 250 mM imidazole in PTB. All FG-N buffers also included 7M guanidine hydrochloride (GuHCl) to prevent aggregation.

### Aggregation time courses

Stocks of PEG (avg. MW 20 kDa, Sigma, Bio-Ultra) and PVP (avg. MW 40 kDa, Sigma) in PTB were prepared at 20 or 40% w/v; L-serine (EMD Millipore Calbiochem) stocks were prepared at 30% w/v. PEG and serine stocks had pH 7; PVP stocks had pH 7 or pH 5 (no difference was found in FG-N aggregation in buffer at pH 5-8). Lyophilized lysate was prepared by homogenizing and centrifuging BL21 DE3 Gold cells. The supernatant was lyophilized in 25 mM NH_4_HCO_3_ and resuspended in PTB immediately before use. A 10 mM stock solution of Thioflavin T (ThT) in PTB was prepared and filtered no more than a week before the time course, stored at room temperature and protected from light.

No more than an hour before beginning recording the ThT fluorescence intensity for the time course, FG-N was desalted into PTB to remove imidazole and GuHCl, and then immediately added to pre-prepared wells containing the appropriate crowding agent if appropriate, 200 *μ*M ThT, and recording was begun. All samples in a given time course were prepared from the same FG-N stock. In each experiment, a sample containing FG-N and 7M GuHCl was prepared as a negative control. For each condition, background samples were prepared containing crowding agent and ThT but without FG-N.

The final sample volume was 150-*μ*L, and experiments were performed in black, flat-bottomed, clear-bottomed 96-well plates. Each well contained a 3mm-diameter glass or Teflon bead [35]. Each sample yielded four to six replicates. A single blank sample was used per condition. The plate was sealed with a PCR seal. The fluorescence was measured at 10-minute intervals with an excitation wavelength of 450 nm, emission wavelength of 482 nm, and 5 nm bandwidth using a Safire II plate reader. The plate shook orbitally at high speed between measurements and was held at a temperature of 30^*°*^C.

In parallel with the sample preparation, the concentration of the desalted FG-N stock was measured with a BCA assay before aggregation could occur. In previous work, [15], we confirmed that the BCA assay gave a reliable measurement of protein concentration, as opposed to a Bradford assay, which we found unreliable. In order to ensure that we caught the early stages of aggregation, we began each experiment before the BCA assay was complete, and so were not able to control the protein concentration. As a result, there was some variability in protein concentration, ranged from 0.3-2.5 mg/mL FG-N. No dependence of aggregation parameters upon FG-N concentration was observed (Fig. S5).

### Aggregation kinetics analysis

The fluorescence intensity of each time course sample was normalized and fit to a sigmoid function in order to extract aggregation rates and lag times. The data were first background-subtracted by subtracting the mean blank intensity. In the cases where the blank intensity changed measurably over the time course, the blank intensity was subtracted pointwise from the sample data. The background-subtracted curves were then fit to a sigmoid given by

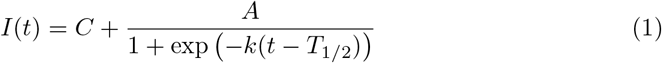

where *I*(*t*) is the normalized fluorescence intensity and *t* is the time since desalting. The aggregation kinetics are described by the rate *k* and the half-time *T*_1*/*2_, which reflects the time needed for the intensity to reach half of its asymptotic value. More descriptive than the half-time is the lag time *T*_*𝓁*_, calculated as

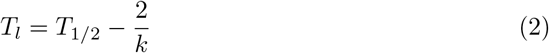

and representing the duration of the lag phase [36]. The lag time here is defined as the intersection of the tangent line of maximum slope of the growth phase with the flat background signal of the lag phase. The remaining fit parameters are an offset *C* and amplitude *A*, which are not used in further analysis.

To account for the unavoidable variations in FG-N concentration in the samples (as final FG-N concentration varied between 0.3 and 2.5 mg/mL), we averaged the fit parameters for all replicates of the FG-N buffer sample within a time course and subtracted that average from all other results. Results from all time courses were then averaged.

In total, for buffer, serine, 13% PEG and 13% PVP, there were 5 different experiments run from 4 biological replicates (different cell pellets and purifications), each with typically 6 replicates. There were five total replicates of the GuHCl condition (4 biological replicates). There were three independent experiments from different biological replicates for lysate. We varied PEG and PVP concentrations in two independent experiments, from different cell pellets and purifications, with with three replicates in each experiment. Error bars on the plots are the standard error of the mean, defined as the standard deviation/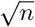, with *n* being the number of replicates. All raw data are available here: https://github.com/lorenhough/FG-N-aggregation

### Viscosity measurements

Dynamic viscosity of PVP and PEG solutions were measured using an Ostwald viscometer with a capillary diameter of 0.5 mm and an efflux volume of 2 mL. Viscometer temperature was maintained at 30^*°*^C by a water bath. Efflux times measured in triplicate for PVP and PEG solutions were 913.4 *±* 0.5 s and 1957 *±* 0.8 s (*±* here refers to standard deviation), respectively, and converted to viscosity based on the efflux times for S3 and S20 certified viscosity standards and the solution densities. Briefly, viscosity for each solution or standard was measured relative to pure water using *η* = *ηω · ρ · t/*(*ρω · tω*), where *ηω* and *ρω* are the known values of viscosity and density, respectively, at 30^*°*^C for pure water and t is the mean efflux time (*tω* for water). Measured viscosity for the S3 and S20 standards was used to estimate error and calibrate the viscometer measurements. Percent error for the S3 and S20 standards, calculated as |measured–certified value|/certified value, was found to be 1.9% and 1.5%, respectively, suggesting that the percent error in the calibrated measurement of viscosity for the PVP and PEG solutions is *<* 2%.

### Fluorescence emission spectra

Samples were prepared by desalting 130 *μ*L of 520 *μ*M FG-N with a Zeba spin desalting column immediately before use to remove the GuHCl. Samples used in fluorescence emission spectra contained 340 or 170 *μ*M FG-N in PTB with 5% w/v PEG, 5% w/v PVP, or no crowder. High concentrations of crowder were not possible due to background fluorescence.

Fluorescence spectra were recorded using a Photon Technology International QM-6 fluorimeter with 4 nm excitation and emission slits and an excitation wavelength of 240 nm. Emission spectra were recorded at 1 nm intervals and automatically averaged over two runs. The PMT detector was set to 1000 V. The emission spectra of background samples containing crowding agents but no protein were also recorded, as well as a buffer-only spectrum. A micro quartz UV cuvette with approximately 50 *μ*L sample volume was used for all spectra. Between runs, the cuvette was rinsed three times with ethanol and five times with deionized water, and the exterior gently blotted with ethanol. Following samples containing aggregated FG-N, the cuvette was rinsed with 7M GuHCl, soaked in 7M GuHCl for five minutes, and cleaned as described above. FG-N samples were tested within two hours of GuHCl removal to avoid any aggregation. The samples were then allowed to incubate at room temperature overnight without shaking and tested again. FG-N samples were visibly cloudy after overnight incubation.

Emission spectra were background-subtracted by averaging over two separate spectra and subtracting the appropriate blank.

### NMR spectroscopy

The NMR samples contained 13% w/v of either PEG or PVP, 140 *μ*M FSFG-K ^15^N, 10% D_2_O, 1% 15 mM TSP or 1% 1mM DSS in PTB. An identical sample containing no crowding agents was also prepared.

NMR experiments were run on an Inova 600 MHz at 25 C. Using the standard proton-detected ^15^N-HSQC experiment from the Varian BioPac. Experiments were taken at 25C, with 128 increments in the indirect dimension, 16 transients per scan, a sweep width of 30 ppm in the nitrogen dimension, and were interleaved. The parameter `relaxT` was adjusted for the measurement of *T*_1_ and *T*_2_: For measurement of *T*_1_, `relaxT` was arrayed (0, 0.1, 0.2, 0.3, 0.4, 0.5, 0.6, 0.7, 0.8, 0.9 ms). For the measurement of *T*_2_, `relaxT` was arrayed (0.01, 0.03, 0.05, 0.07, 0.09, 0.11, 0.13, 0.15, 0.17, 0.19, 0.21, 0.23, 0.25 ms). The data were processed using standard scripts in NMRPipe, and analyzed using CCPNmr Analysis software and depositied in the BMRB: doi:10.13018/BMR50553

## Results and discussion

To probe the aggregation kinetics of FG-N, we recorded aggregation time courses in the presence of multiple crowders, including inert crowders as well as nonspecifically binding ones, and thioflavin T (ThT), a dye which is sensitive to amyloid formation [27, 28]. PEG and PVP were chosen as representative inert crowding agents. Cell lysate was used to imitate the cellular environment and test the effect of nonspecific interactions on FG-N aggregation. Finally, serine was used as a crowder as previous work demonstrated that it increased the aggregation rate of Nup153 [5]. Over the course of several hours, we observed increased fluorescence intensity from ThT concurrent with FG-N aggregation. Increases in ThT fluorescence are typically attributed to amyloid formation, but can result from interaction with other biomacromolecules, and other beta-strand-rich structures [49, 50]. However, given that FG-N has been previously been shown to form amyloid-like structures, we attribute this rise in fluorescence to the formation of amyloid-like aggregates of FG-N [22]. The resulting fluorescence intensity traces showed a lag phase typical of a nucleation phase, followed by a growth phase of rapid aggregation, ending in a plateau phase (Fig. 2A). The plateau phase in some experiments showed a subsequent decrease in fluoresence over time, the origins of which are unknown. The lag time *T*_*𝓁*_ and aggregation rate *k* for each crowding condition were calculated by fitting the intensity to a sigmoid (see Materials and methods).

**Figure 2.**
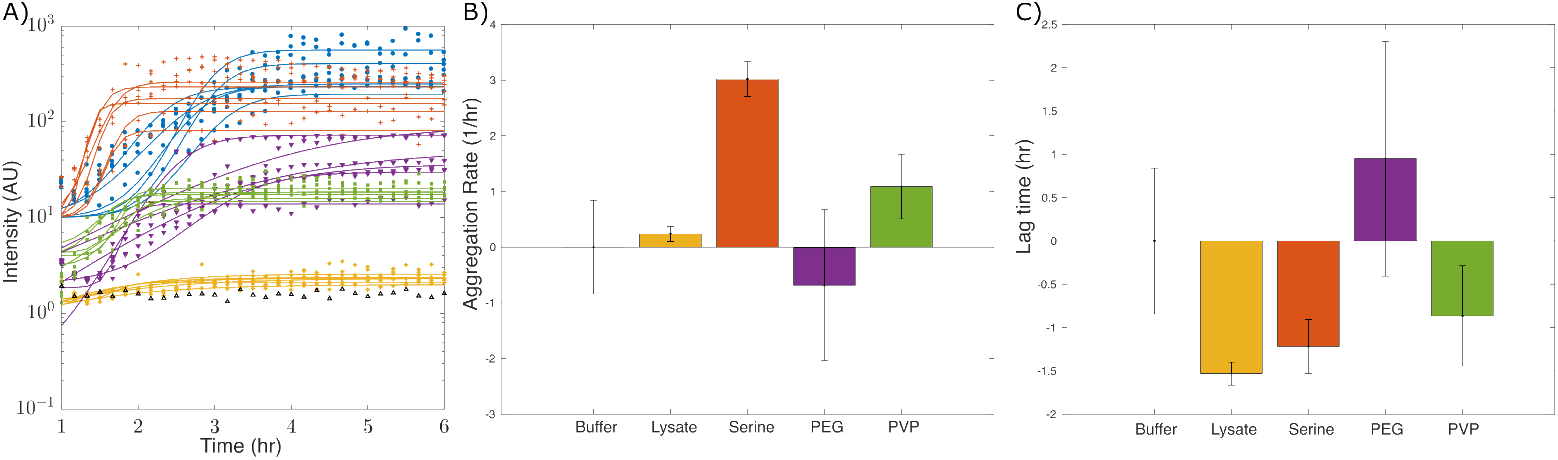
Aggregation kinetics assays. A: Background-subtracted intensity as a function of time for all replicates of a single time course with a single FG-N concentration. Sigmoidal fits are shown (see Materials and Methods). Crowding agent concentrations were 19% w/v serine (orange), 13% PEG (purple), 13% PVP (PVP), and approximately 10 mg/mL lysate (yellow). The buffer condition did not contain a crowding agent. Guanidine hydrochloride concentration was 7M. B: Relative aggregation rates *k* and C: lag times *T*_*𝓁*_ for all crowding conditions. For each independent time course, we subtracted the no-crowding (buffer) condition to obtain the relative aggregation rate and lag times. We then averaged these relative values over all replicates. Error bars are standard error of the mean.

There were changes in both lag time and aggregation rate of FG-N as a result of changing crowder conditions. For both the aggregation rate *k* and lag time *T*_*𝓁*_, a one-factor ANOVA rejected at the *p* = 0.05 level the null hypothesis that there was no difference between the condition means (Fig. 2B, C). We then ran two-tailed Student’s t-tests comparing all conditions. The resulting *p*-values were below 0.05 except for those comparing no crowder to PVP (rate), no crowder to PEG (lag time), and serine to PVP (lag time).

Two crowding agents, PEG and PVP, warranted special attention, as these crowders are commonly considered inert and used interchangeably, but in our experiments, their affects on aggregation kinetics differ. The presence of 13% w/v PEG reduces the aggregation rate *k* as compared to aggregation in buffer, but the same PVP concentration has no measurable effect on the aggregation rate. In contrast, the PVP sample has a significantly shorter lag time *T*_*𝓁*_ than the no-crowder condition, but the PEG sample does not. In order to investigate any differences between the effect of PEG and PVP on aggregation, time courses were run with varying PEG and PVP concentrations (Fig. 3). No trends are apparent in the rate *k* as the concentration of crowding agent changes. Lag time *T*_*𝓁*_ generally decreases with decreasing crowding for both PEG and PVP.

**Figure 3.**
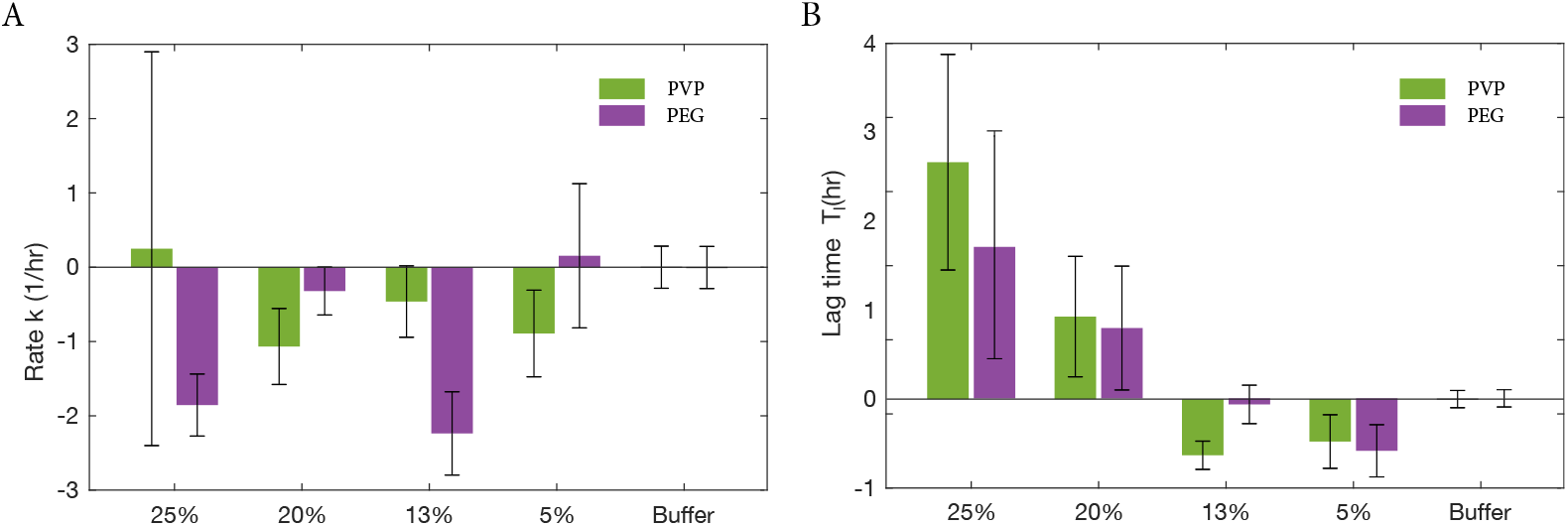
Aggregation kinetics at varying inert crowder concentrations. A: Aggregation rates *k* and B: lag times *T*_*𝓁*_ for varying PEG (purple) and PVP (green) concentrations, rescaled to the no-crowding condition and averaged over all time courses and replicates. Error bars are standard error of the mean.

A possible source of the differences between PEG and PVP as crowding agents is their differing viscosity. A 13% PEG solution in the buffer we named PTB (see methods) has a dynamic viscosity of 15.87 mPa*·*s, while that of a 13% PVP solution in PTB is 7.34 mPa*·*s. Increasing viscosity increases the lag time and decreases the aggregation rate of insulin and *α* synuclein [7, 8]. However, other work suggests that the behavior is non-monotonic depending on viscosity and the aggregation propensity of the protein [4, 30]. The trends we observe are consistent with a non-monotonic dependence on viscosity, in which crowding effects decrease lag time at moderate crowder concentrations but have the opposite effect as viscosity effects begin to dominate.

Another possibility is that the presence of an aromatic ring in PVP, and its interaction with FG peptides, was responsible for the differences in aggregation kinetics between PEG and PVP samples. PVP possesses an aromatic ring, as do the phenylalanine residues of FG-N, while PEG does not. Ring-stacking interactions may therefore play a role in FG-N aggregation in the presence of PVP. Moreover, phenylalanine residues are key to the function of the NPC. While phenylalanine fluorescence is less commonly studied than that of tryptophan, we reasoned that changes in the chemical microenvironment of phenylalanine would result in changes in the fluorescence spectra as is seen for tryptophan [31, 32]. Phenylalanine fluorescence is known to be sensitive to the local environment [29], and has previously been used as probe for amyloid formation due to changes in fluorescence spectra attributed to *π − π\** stacking [47, 48]. Fortuitously, our samples lack either tryptophan and tyrosine residues, making the phenylalanine fluorescence the strongest fluorescence signal among residue side-chains in the near UV.

We collected fluorescence spectra near the emission maximum of phenylalanine from both fresh and aggregated FG-N in either buffer alone, or with 5 % w/v PEG or PVP (Fig. 4). The PEG sample showed the largest difference in peak height and location upon aggregation, with the PVP sample showing the smallest changes. We hypothesized that the phenylalanine residues are in a more similar environment when comparing PVP and aggregated samples than PEG and aggregated samples because ring-stacking interactions are already available in PVP. Both samples with crowding agents developed a peak upon aggregation centered near 310 nm not apparent in the aggregated sample without crowding agent, which may indicate a different structure of the aggregates, though we did not pursue this further (Figs. 4, S7). This is consistent with previous reports of the appearance of a peak at 303-313 nm upon amyloid fiber formation of an amyloid peptide fragment, and has been attributed to *π − π\** stacking of the phenyl units in amyloids [47, 48]. The broad shoulder at longer wavelengths (320-340 nm) may arise from more dynamic interactions between phenylalanine residues, or between phenylalnine residues and the ring of PVP, as a peak centered at 320 was previously found at high concentration of phenylalanine in solution and attributed to dynamic interactions between phenyl rings. [46]

**Figure 4.**
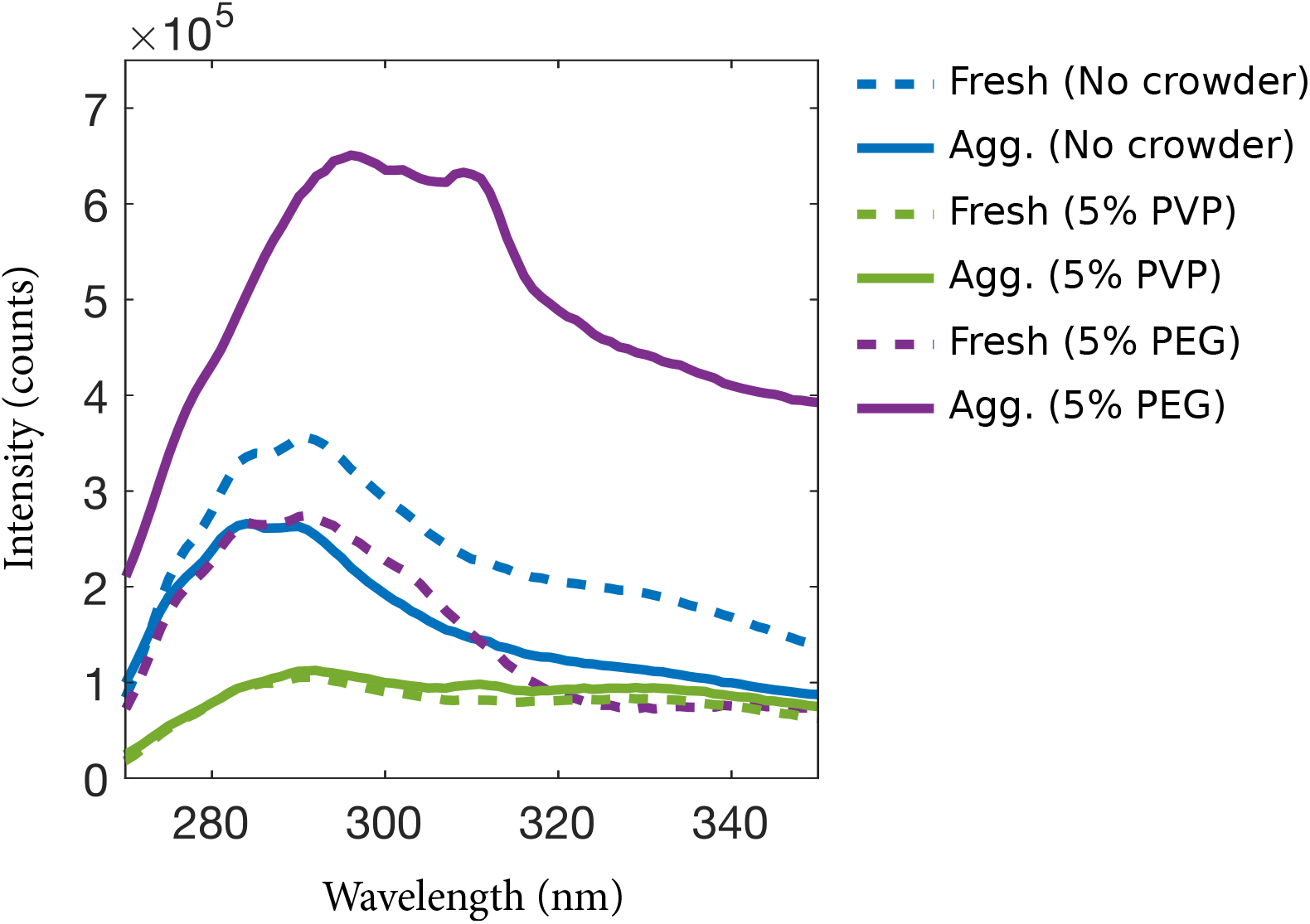
Fluorescence spectra of fresh and aggregated FG-N in 5% PEG, 5% PVP, and without a crowding agent. All data were background-subtracted.

In analogy to tryptophan fluorescence shifts, these results suggest that the chemical microenvironment of the phenylalanine residues in FG-N changes upon aggregation under crowded conditions. As a general rule, the emission spectrum of tryptophan becomes redshifted when the tryptophan residue is surrounded by an increasingly hydrophilic environment [31, 32]. It is not clear that a similar shift should hold true in detail for phenylalanine, but the differential shift in the location of the main emission peak between PEG and PVP conditions is consistent with a difference in the chemical microenvironment of the phenylalanine residues. Moreover, the fluorescence emission maximum for FG-K, which is generally more hydrophilic than FG-N, is blue-shifted relative to FG-N (Fig. S7).

To further investigate the effect of crowding agents on the residues of FG Nups, we measured the longitudinal (*R*_1_) and transverse (*R*_2_) relaxation rates of the non-aggregating peptide FSFG-K using NMR spectroscopy. FG-N cannot be used in NMR experiments due to its aggregation propensity which results in loss of signal intensity. Therefore, we used the well-characterized peptide FSFG-K (Fig. 1), which is of a similar length to FG-N and contains a similar number of FG motifs but does not aggregate. We measured the relaxation rates of FSFG-K in 13% w/v PEG or PVP and compared the rates to those of FSFG-K in non-crowded conditions (Fig. 5). Due to the disordered nature and highly repetitive nature of the FG repeats in FSFG-K, many peaks in the NMR spectra overlap. This effect is particularly pronounced for the FSFG motif itself, in which the six repeated motifs are almost entirely collapsed to four peaks, one for each of the residue in ‘FSFG’. The sole exception is the phenylalanine closest to the N-terminal, which has a peak distinct from the others. Therefore, we broke the sequence of the FSFG-K peptide into quasi-repeating segments, each 19 amino acids in length, which begin with an FSFG motif and contain the hydrophilic linker which separates it from the next FSFG motif (Fig. 5B, inset). We then averaged the signals over the first five repeats.

**Figure 5.**
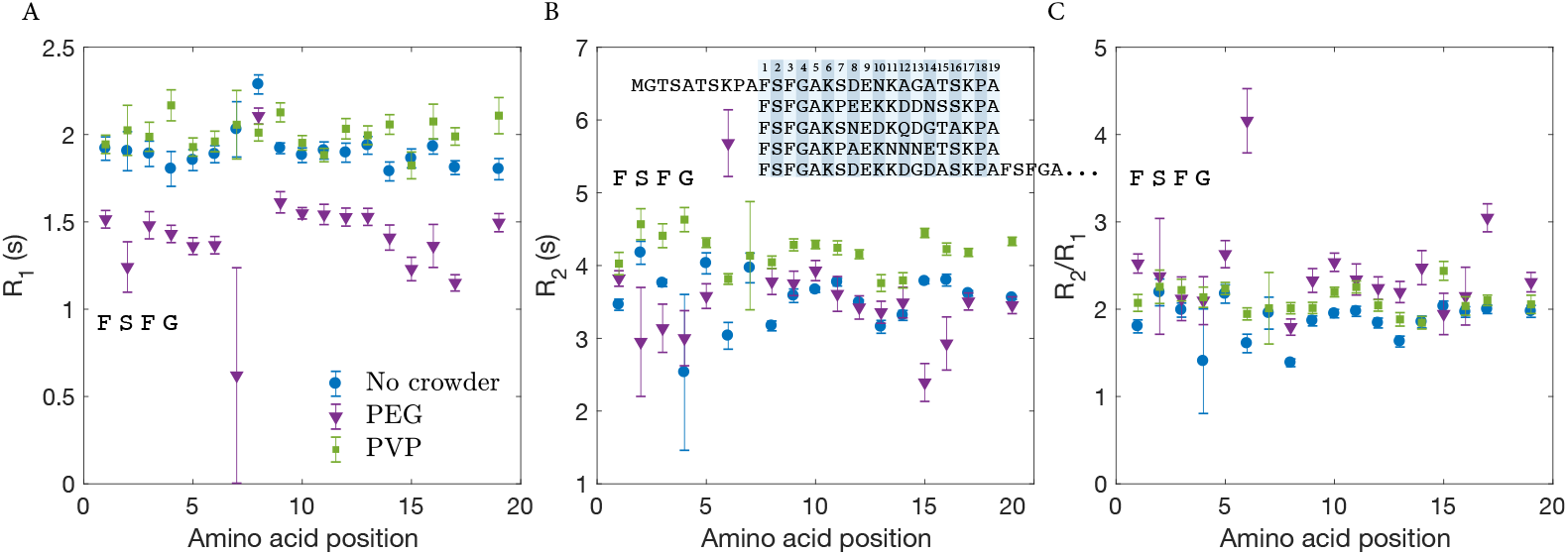
Relaxation rates for FSFG-K in 13% PEG, 13% PVP, or PTB only. The 19-amino-acid quasi-repeating unit that begins with the FSFG motif has been averaged over the first five repeats as shown. A: Longitudinal relaxation rate *R*_1_, B: transverse relaxation rate *R*_2_, C: Ratio *R*_2_*/R*_1_. The errors in *R*_1_ and *R*_2_ are given by the weighted average of the fit errors for the unique peaks that are being averaged [33]. The errors of the ratio *R*_2_*/R*_1_ were propagated from *R*_1_ and *R*_2_ errors.

The crowded conditions show noticeable differences in relaxation rates compared to the buffer condition across the whole peptide. In general, polymeric crowders can impact intrinsically disordered regions through a variety of mechanisms, including structural changes that result in compaction, interactions between the crowding agent and the protein, and through changes in the solution viscosity [37–39, 41, 42, 45]. The systemic differences in rates do not all correspond to changes in viscosity, as is typically seen for solutions made viscous by high concentrations of small molecules [41]. This has also been observed in other systems, where the local dynamics were thought to not be impacted by polymeric crowders [37]. The PVP sample shows the highest *R*_1_ and *R*_2_ values despite having an intermediate viscosity. Likewise, the *R*_1_ values are noticeably lower for in the PEG condition, which is also not easily explained by viscosity changes.

## Conclusion

We used several methods to investigate FG Nup aggregation in crowded conditions, focusing on FG-N as a representative peptide (Fig. 1). Using both aggregation kinetics and NMR specroscopy spin-relaxation measurements, we observed differences in aggregation and the resulting microenvironments of the peptide’s residues depending on the presence and type of crowder. The aggregation timecourses pointed to differences between most of the crowders tested (Fig. 2), and between PEG and PVP in particular (Fig. 3). In order to further determine the differences between PEG and PVP on FG-N aggregation, we pursued fluorimetry and NMR. Results from both reinforced the idea that there are in fact differences between FG-N behavior in these crowders, consistent with other work highlighting interactions between crowding agents and proteins [1, 51–53]. Intrinsic phenylalanine fluorescence in aggregated FG-N revealed an emission peak at 310 nm that was missing from buffer only conditions, and suggests structural differences for individual residues (Fig. 4). Likewise, NMR measured relaxation rates for residues in FSFG-K showed differences in PEG, PVP, and buffer-only conditions which did not trend with changes in solution viscosity, and thus imply a structural origin. Since FG repeats are involved not only to binding transport factors for selective transport, but also in forming transient crosslinks with other FG Nups, our results indicate that great care should be taken in determining what conditions to use to study Nups *in vitro*. Selecting among even relatively inert crowders, such as PEG and PVP, potentially modulates FG Nup cohesiveness and dynamical behavior.

## Supporting information

**S1 File**. This file contains images of SDS PAGE results showing a typical purification and protein purity, additional plots of aggregation timecourses, buffer controls lacking proteins, disaggregation of the data by date and protein concentration, fluoresence spectra of FG-N normalized by total intensity, and of FG-K for comparison, and the NMR relaxation data for each condition as a function of primary sequence.

## Acknowledgments

We thank Meredith Betterton for useful discussions. Fluorescence experiments were performed in the Shared Instruments Pool (RRID: SCR 018986) in the Department of Biochemistry at the University of Colorado Boulder

## References

1. Breydo L, Reddy KD, Piai A, Felli IC, Pierattelli R, Uversky VN. The Crowd You’re in with: Effects of Different Types of Crowding Agents on Protein Aggregation. Biochimica et Biophysica Acta (BBA) - Proteins and Proteomics. 2014;1844(2):346–357. doi:10.1016/j.bbapap.2013.11.004.

2. Breydo L, Sales AE, Frege T, Howell MC, Zaslavsky BY, Uversky VN. Effects of Polymer Hydrophobicity on Protein Structure and Aggregation Kinetics in Crowded Milieu. Biochemistry. 2015;54(19):2957–2966. doi:10.1021/acs.biochem.5b00116.

3. Lee CF, Bird S, Shaw M, Jean L, Vaux DJ. Combined Effects of Agitation, Macromolecular Crowding, and Interfaces on Amyloidogenesis. Journal of Biological Chemistry. 2012;287(45):38006–38019. doi:10.1074/jbc.M112.400580.

4. Magno A, Caflisch A, Pellarin R. Crowding Effects on Amyloid Aggregation Kinetics. The Journal of Physical Chemistry Letters. 2010;1(20):3027–3032. doi:10.1021/jz100967z.

5. Milles S, Huy Bui K, Koehler C, Eltsov M, Beck M, Lemke EA. Facilitated Aggregation of FG Nucleoporins under Molecular Crowding Conditions. EMBO reports. 2013;14(2):178–183. doi:10.1038/embor.2012.204.

6. Sukenik, Shahar and Harries, Daniel Insights into the Disparate Action of Osmolytes and Macromolecular Crowders on Amyloid Formation Prion. 2012; 6:26–31. doi:10.4161/pri.6.1.18132

7. Sleutel M, Van Driessche AES, Pan W, Reichel EK, Maes D, Vekilov PG. Does Solution Viscosity Scale the Rate of Aggregation of Folded Proteins? The Journal of Physical Chemistry Letters. 2012;3(10):1258–1263. doi:10.1021/jz300459n.

8. Saha S, Sharma A, Deep S. Differential Influence of Additives on the Various Stages of Insulin Aggregation. RSC Advances. 2016;6(34):28640–28652. doi:10.1039/C5RA27206H.

9. Cheng, Xian and Shkel, Irina A. and O′Connor, Kevin and Record, M. Thomas Experimentally Determined Strengths of Favorable and Unfavorable Interactions of Amide Atoms Involved in Protein Self-Assembly in Water. Proceedings of the National Academy of Sciences. 2020. doi:10.1073/pnas.2012481117.

10. Shkel, Irina A. and Knowles, D. B. and Record, M. Thomas Separating Chemical and Excluded Volume Interactions of Polyethylene Glycols with Native Proteins: Comparison with PEG Effects on DNA Helix Formation Biopolymers. 2015; 105:517–527. doi:10.1002/bip.22662

11. Chang, Yu-Chu and Oas, Terrence G. Osmolyte-Induced Folding of an Intrinsically Disordered Protein: Folding Mechanism in the Absence of Ligand Biochemistry. 2010; 49:5086–5096/ doi:10.1021/bi100222h

12. Protter DSW, Rao BS, Treeck BV, Lin Y, Mizoue L, Rosen MK, et al. Intrinsically Disordered Regions Can Contribute Promiscuous Interactions to RNP Granule Assembly. Cell Reports. 2018;22(6):1401–1412. doi:10.1016/j.celrep.2018.01.036.

13. Jovanovic-Talisman T, Zilman A. Protein Transport by the Nuclear Pore Complex: Simple Biophysics of a Complex Biomachine. Biophysical Journal. 2017;113(1):6–14. doi:10.1016/j.bpj.2017.05.024.

14. Timney BL, Raveh B, Mironska R, Trivedi JM, Kim SJ, Russel D, et al. Simple Rules for Passive Diffusion through the Nuclear Pore Complex. J Cell Biol. 2016; p. jcb.201601004. doi:10.1083/jcb.201601004.

15. Hough LE, Dutta K, Sparks S, Temel DB, Kamal A, Tetenbaum-Novatt J, et al. The Molecular Mechanism of Nuclear Transport Revealed by Atomic Scale Measurements. eLife. 2015; p. e10027. doi:10.7554/eLife.10027.

16. Gu C, Vovk A, Zheng T, Coalson RD, Zilman A. The Role of Cohesiveness in the Permeability of the Spatial Assemblies of FG Nucleoporins. Biophysical Journal. 2019;116(7):1204–1215. doi:10.1016/j.bpj.2019.02.028.

17. Tagliazucchi M, Peleg O, Kroeger M, Rabin Y, Szleifer I. Effect of Charge, Hydrophobicity, and Sequence of Nucleoporins on the Translocation of Model Particles through the Nuclear Pore Complex. Proceedings of the National Academy of Sciences of the United States of America. 2013;110(9):3363–3368. doi:10.1073/pnas.1212909110.

18. Nasrabad AE, Jasnow D, Zilman A, Coalson RD. Precise Control of Polymer Coated Nanopores by Nanoparticle Additives: Insights from Computational Modeling. Journal of Chemical Physics. 2016;145(6):064901. doi:10.1063/1.4955191.

19. Mincer JS, Simon SM. Simulations of Nuclear Pore Transport Yield Mechanistic Insights and Quantitative Predictions. Proceedings Of The National Academy Of Sciences Of The United States Of America. 2011;108(31):E351–8. doi:10.1073/pnas.1104521108.

20. Frey S, Richter R, Görlich D. FG-Rich Repeats of Nuclear Pore Proteins Form a Three-Dimensional Meshwork with Hydrogel-like Properties. Science. 2006;314(5800):815–817.

21. Frey S, Görlich D. A Saturated FG-Repeat Hydrogel Can Reproduce the Permeability Properties of Nuclear Pore Complexes. Cell. 2007;130(3):512–523. doi:10.1016/j.cell.2007.06.024.

22. Ader C, Frey S, Maas W, Schmidt HB, Gorlich D, Baldus M. Amyloid-like Interactions within Nucleoporin FG Hydrogels. Proceedings of the National Academy of Sciences. 2010;107(14):6281–6285. doi:10.1073/pnas.0910163107.

23. Schmidt HB, Görlich D. Nup98 FG Domains from Diverse Species Spontaneously Phase-Separate into Particles with Nuclear Pore-like Permselectivity. eLife. 2015;4:e04251. doi:10.7554/eLife.04251.

24. Jahn TR, Radford SE. Folding versus Aggregation: Polypeptide Conformations on Competing Pathways. Archives of Biochemistry and Biophysics. 2008;469(1):100–117. doi:10.1016/j.abb.2007.05.015.

25. Sakiyama Y, Mazur A, Kapinos LE, Lim RYH. Spatiotemporal Dynamics of the Nuclear Pore Complex Transport Barrier Resolved by High-Speed Atomic Force Microscopy. Nature Nanotechnology. 2016;advance online publication. doi:10.1038/nnano.2016.62.

26. Moussavi-Baygi R, Mofrad MRK. Rapid Brownian Motion Primes Ultrafast Reconstruction of Intrinsically Disordered Phe-Gly Repeats Inside the Nuclear Pore Complex. Scientific Reports. 2016;6. doi:10.1038/srep29991.

27. Picken MM, Herrera GA. Thioflavin T Stain: An Easier and More Sensitive Method for Amyloid Detection. In: Fasn Mmpm, M D AD, M D Gah, editors. Amyloid and Related Disorders. Current Clinical Pathology. Humana Press; 2012. p. 187–189.

28. Groenning M. Binding Mode of Thioflavin T and Other Molecular Probes in the Context of Amyloid Fibrils—Current Status. Journal of Chemical Biology. 2009;3(1):1–18. doi:10.1007/s12154-009-0027-5.

29. Sudhakar, K., W.W. Wright, S.A. Williams, C.M. Phillips, and J.M. Vanderkooi. Phenylalanine fluorescence and phosphorescence used as a probe of conformation for cod parvalbumin. J Fluoresc. 1993;3:57–64. doi:10.1007/BF00865318.

30. Munishkina LA, Cooper EM, Uversky VN, Fink AL. The Effect of Macromolecular Crowding on Protein Aggregation and Amyloid Fibril Formation. Journal of Molecular Recognition. 2004;17(5):456–464. doi:10.1002/jmr.699.

31. Ladokhin AS. Fluorescence Spectroscopy in Peptide and Protein Analysis. In: Meyers RA, editor. Encyclopedia of Analytical Chemistry. Chichester, UK: John Wiley & Sons, Ltd; 2000. p. a1611.

32. Serrano-Andrés L, Roos BO. Theoretical Study of the Absorption and Emission Spectra of Indole in the Gas Phase and in a Solvent. Journal of the American Chemical Society. 1996;118(1):185–195. doi:10.1021/ja952035i.

33. Taylor J. An Introduction to Error Analysis. 2nd ed. University Science Books; 1997.

34. Magnusdottir A, Johansson I, Dahlgren LG, Nordlund P, Berglund H. Enabling IMAC Purification of Low Abundance Recombinant Proteins from E. Coli Lysates. Nature Publishing Group. 2009;6(7):477–478. doi:10.1038/nmeth0709-477.

35. Giehm L, Otzen DE. Strategies to Increase the Reproducibility of Protein Fibrillization in Plate Reader Assays. Analytical Biochemistry. 2010;400(2):270–281. doi:10.1016/j.ab.2010.02.001.

36. Arosio P, Knowles TPJ, Linse S. On the Lag Phase in Amyloid Fibril Formation. Physical Chemistry Chemical Physics. 2015;17(12):7606–7618. doi:10.1039/c4cp05563b.

37. Cino, E., Karttunen, M. & Choy, W. Effects of Molecular Crowding on the Dynamics of Intrinsically Disordered Proteins. PLOS ONE. 7, e49876 (2012,11), https://journals.plos.org/plosone/article?id=10.1371/journal.pone.0049876, Publisher: Public Library of Science

38. Fonin, A., Darling, A., Kuznetsova, I., Turoverov, K. & Uversky, V. Intrinsically disordered proteins in crowded milieu: when chaos prevails within the cellular gumbo. Cellular And Molecular Life Sciences. 75, 3907–3929 (2018,11), 10.1007/s00018-018-2894-9

39. Bonucci, A., Palomino-Schätzlein, M., Molina, P., Arbe, A., Pierattelli, R., Rizzuti, B., Iovanna, J. & Neira, J. Crowding Effects on the Structure and Dynamics of the Intrinsically Disordered Nuclear Chromatin Protein NUPR1. Frontiers In Molecular Biosciences. 8 (2021), https://www.frontiersin.org/articles/10.3389/fmolb.2021.684622

40. Hu, Y., Cheng, K., He, L., Zhang, X., Jiang, B., Jiang, L., Li, C., Wang, G., Yang, Y. & Liu, M. NMR-Based Methods for Protein Analysis. Analytical Chemistry. 93, 1866–1879 (2021,2), 10.1021/acs.analchem.0c03830, Publisher: American Chemical Society

41. Szasz, C., Alexa, A., Toth, K., Rakacs, M., Langowski, J. & Tompa, P. Protein Disorder Prevails under Crowded Conditions. Biochemistry. 50, 5834–5844 (2011,7), 10.1021/bi200365j, Publisher: American Chemical Society

42. Shahid, S., Hassan, M., Islam, A. & Ahmad, F. Size-dependent studies of macromolecular crowding on the thermodynamic stability, structure and functional activity of proteins: in vitro and in silico approaches. Biochimica Et Biophysica Acta (BBA) - General Subjects. 1861, 178–197 (2017,2), https://www.sciencedirect.com/science/article/pii/S0304416516304226

43. Grudziaž, K., Zawadzka-Kazimierczuk, A. & Kózmiński, W. High-dimensional NMR methods for intrinsically disordered proteins studies. Methods. 148 pp. 81–87 (2018,9), https://www.sciencedirect.com/science/article/pii/S1046202318300082

44. Hu, Y., Cheng, K., He, L., Zhang, X., Jiang, B., Jiang, L., Li, C., Wang, G., Yang, Y. & Liu, M. NMR-Based Methods for Protein Analysis. Analytical Chemistry. 93, 1866–1879 (2021,2), 10.1021/acs.analchem.0c03830, Publisher: American Chemical Society

45. Adamski, W., Salvi, N., Maurin, D., Magnat, J., Milles, S., Jensen, M., Abyzov, A., Moreau, C. & Blackledge, M. A Unified Description of Intrinsically Disordered Protein Dynamics under Physiological Conditions Using NMR Spectroscopy. Journal Of The American Chemical Society. 141, 17817–17829 (2019,11), 10.1021/jacs.9b09002, Publisher: American Chemical Society

46. Leroy, E., H. Lami, and G. Laustriat. Fluorescence Lifetime and Quantum Yield of Phenylalanine Aqueous Solutions. Temperature and Concentration Effects. Photochemistry and Photobiology. 13, 411–421 (1971), 10.1111/j.1751-1097.1971.tb06132.x, Publisher: Wiley Online Library

47. Krysmann, M.J., V. Castelletto, A. Kelarakis, I.W. Hamley, R.A. Hule, and D.J. Pochan. Self-Assembly and Hydrogelation of an Amyloid Peptide Fragment. Biochemistry. 47, 4597–4605 (2008), 10.1021/bi8000616, Publisher: American Chemical Society

48. Marshall, K.E., K.L. Morris, D. Charlton, N. O’Reilly, L. Lewis, H. Walden, and L.C. Serpell. Hydrophobic, Aromatic, and Electrostatic Interactions Play a Central Role in Amyloid Fibril Formation and Stability. Biochemistry. 50, 2061–2071 (2011), 10.1021/bi101936c, Publisher: American Chemical Society

49. Gade Malmos, K., L.M. Blancas-Mejia, B. Weber, J. Buchner, M. Ramirez-Alvarado, H. Naiki, and D. Otzen. ThT 101: a primer on the use of thioflavin T to investigate amyloid formation. Amyloid. 24, 1–16 (2017), 10.1080/13506129.2017.1304905, Publisher: Taylor and Francis

50. Biancalana, M., and S. Koide. Molecular Mechanism of Thioflavin-T Binding to Amyloid Fibrils. Biochim Biophys Acta. 1804, 1405–1412 (2010). 10.1016/j.bbapap.2010.04.001 Publisher: Elsevier

51. Miklos, A.C., C. Li, N.G. Sharaf, and G.J. Pielak. Volume Exclusion and Soft Interaction Effects on Protein Stability under Crowded Conditions. Biochemistry. 49, 6984–6991. (2010). 10.1021/bi100727y, Publisher: American Chemical Society

52. Wu, J., C. Zhao, W. Lin, R. Hu, Q. Wang, H. Chen, L. Li, S. Chen, and J. Zheng. Binding characteristics between polyethylene glycol (PEG) and proteins in aqueous solution. Journal of Materials Chemistry B. 2, 2983–2992 (2014). 10.1039/C4TB00253A Publisher: Royal Society of Chemistry

53. Bekale, L., D. Agudelo, and H.A. Tajmir-Riahi. The role of polymer size and hydrophobic end-group in PEG–protein interaction. Colloids and Surfaces B: Biointerfaces. 130, 141–148 (2015). 10.1016/j.colsurfb.2015.03.045 Publisher: Elsevier

