## Supplemental Figures for "Aggregation of an FG nucleoporin under crowded conditions"

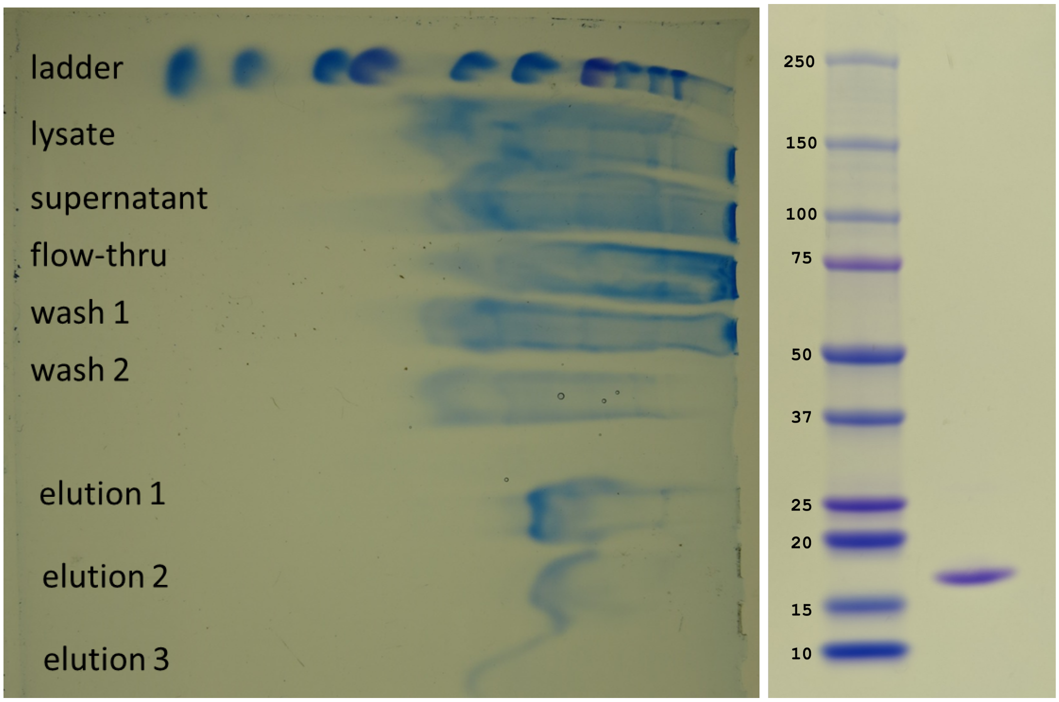


**Figure 1. (left) Example SDS PAGE gel showing the purity of our FG-N protein. The gel is smeared because of the high concentration of guanidine hydrochloride. (right) Desalted protein was rerun to show the sample purity.**

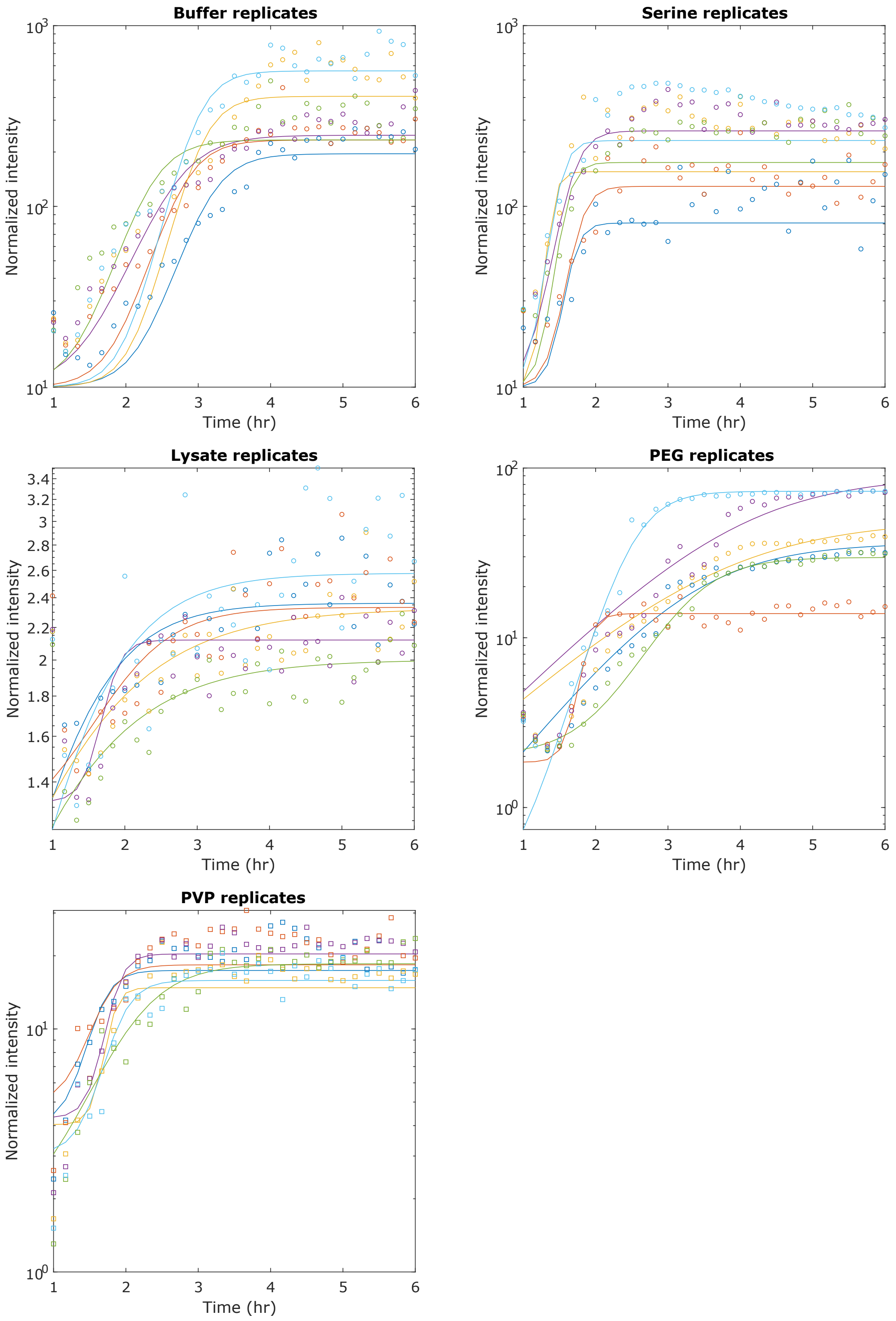




**Figure 2. Normalized thioflavin intensity for individual aggregation replicates for each condition, taken from a single experimental run.** Each is divided by the intensity in a buffer-only well to normalize.

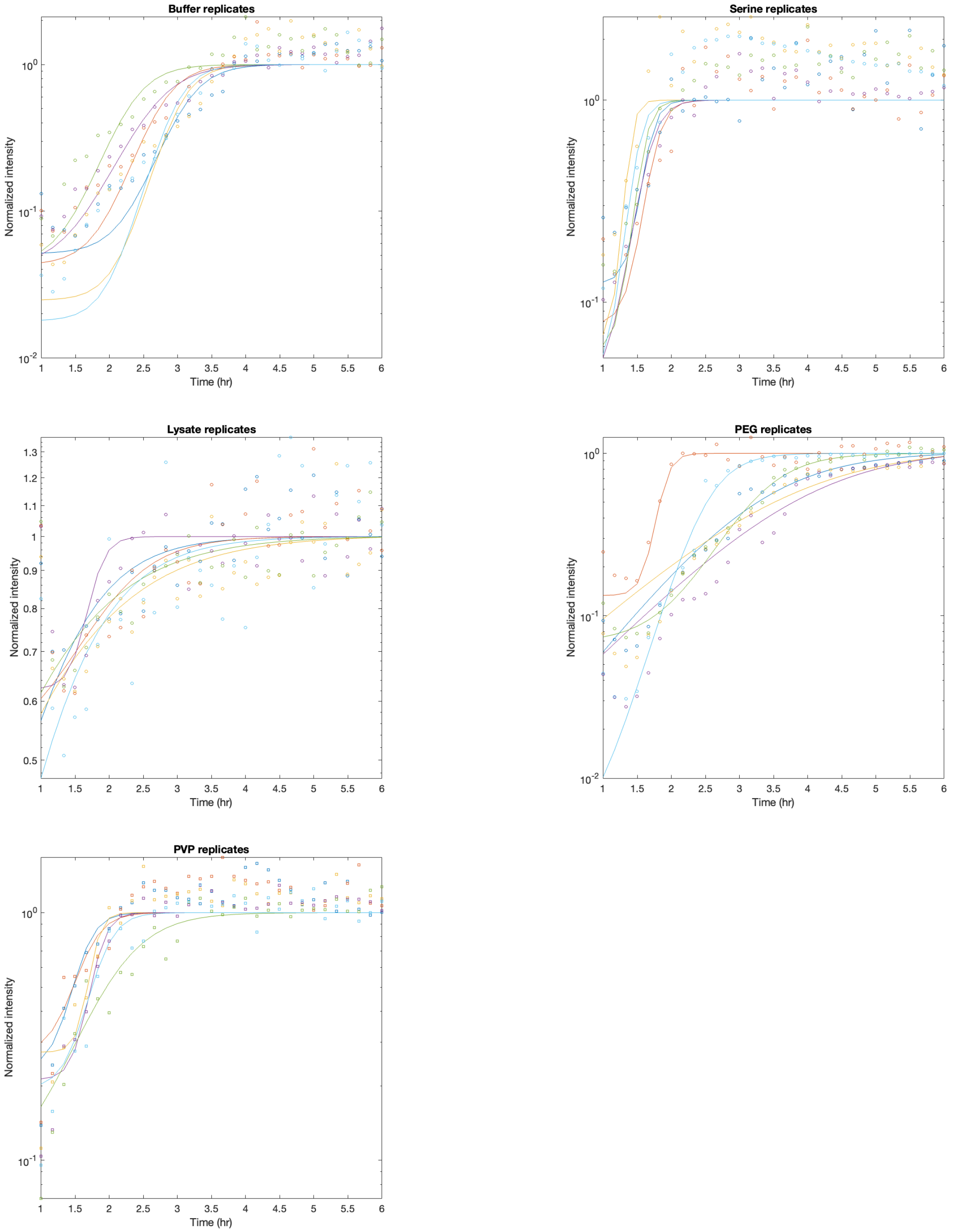


**Figure 3. Normalized thioflavin intensity for individual aggregation replicates for each condition, taken from a single experimental run.** Each is divided by the amplitude from the fit to normalize.

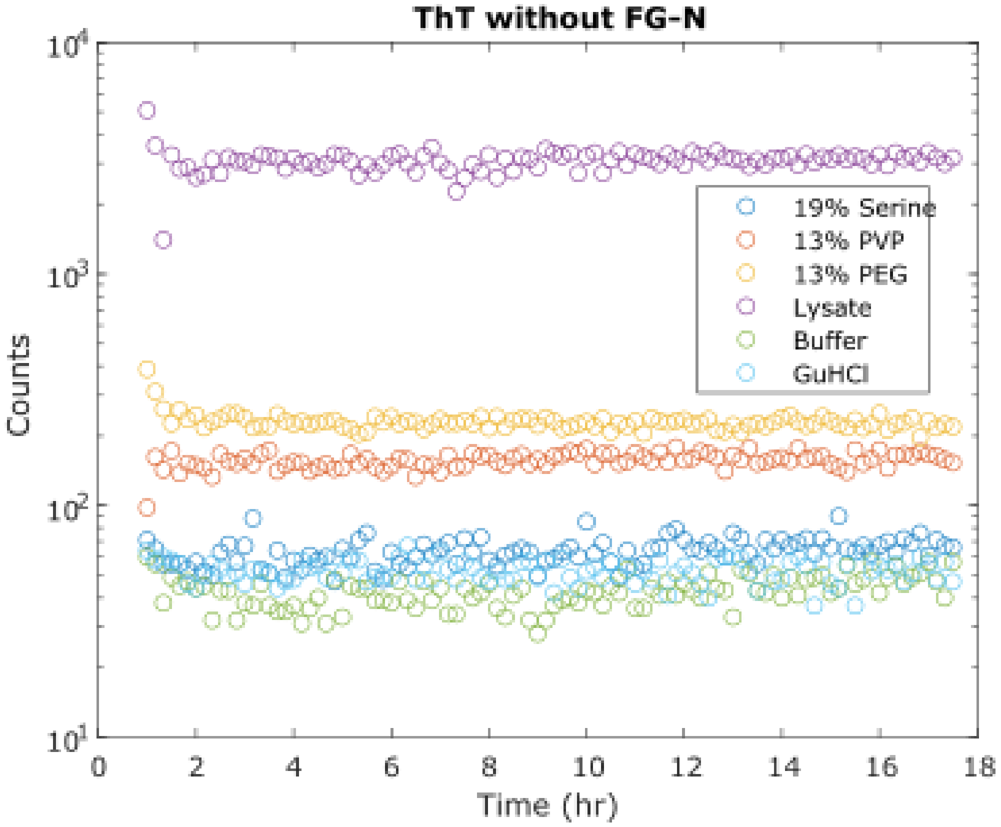


**Figure 4. Thioflavin fluorescent intensity as a function of time for the buffer wells, lacking protein.** Even in the absence of protein, the intensities varied dramatically for different buffer conditions.

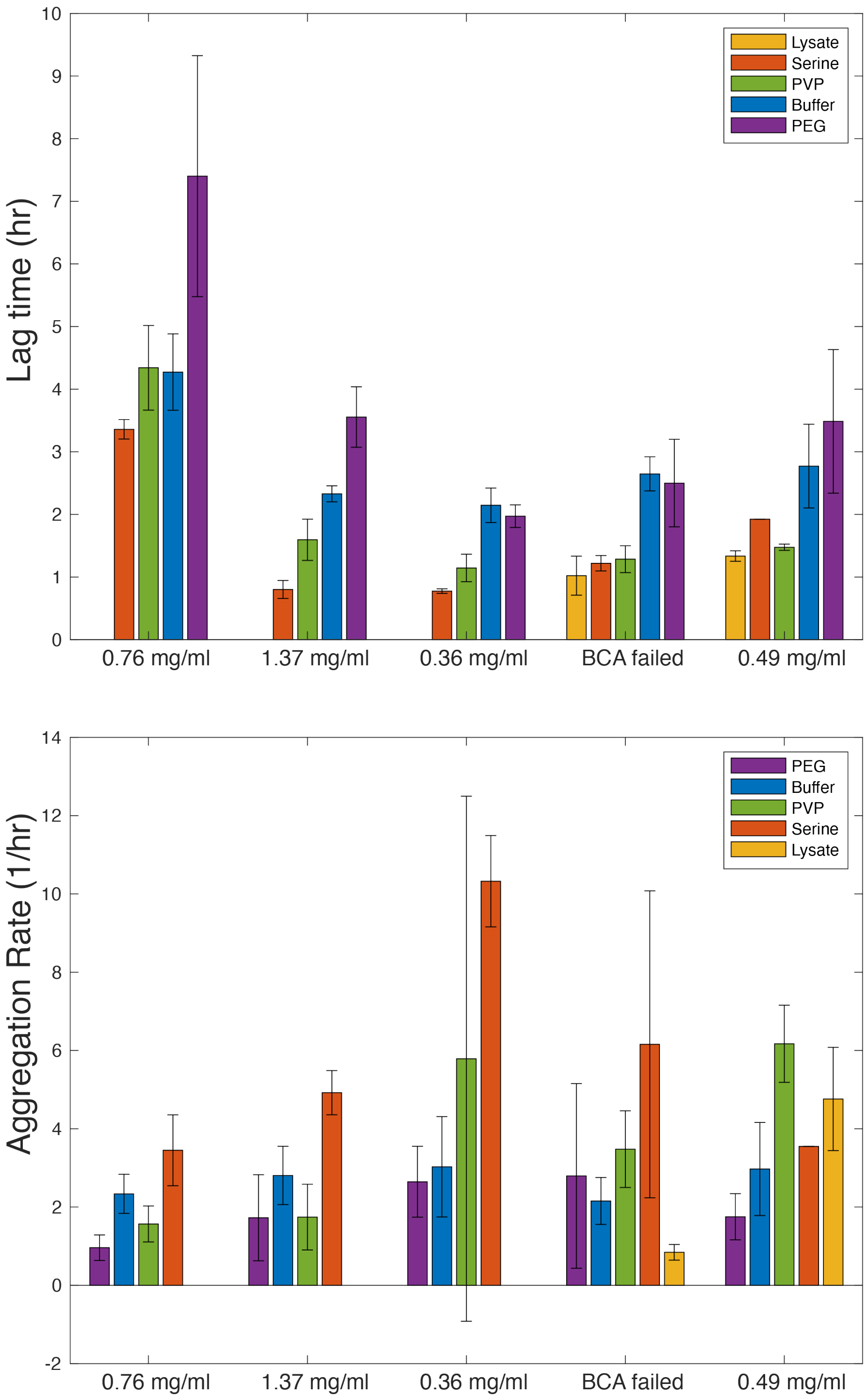


**Figure 5. Lag time (top) and aggregation rate (bottom) vs. protein concentration for each experimental conditions.** Even though aggregation typically shows some concentration dependence, we did not see significant dependence on concentration. The trends in dependence on condition were similar across all experiments.

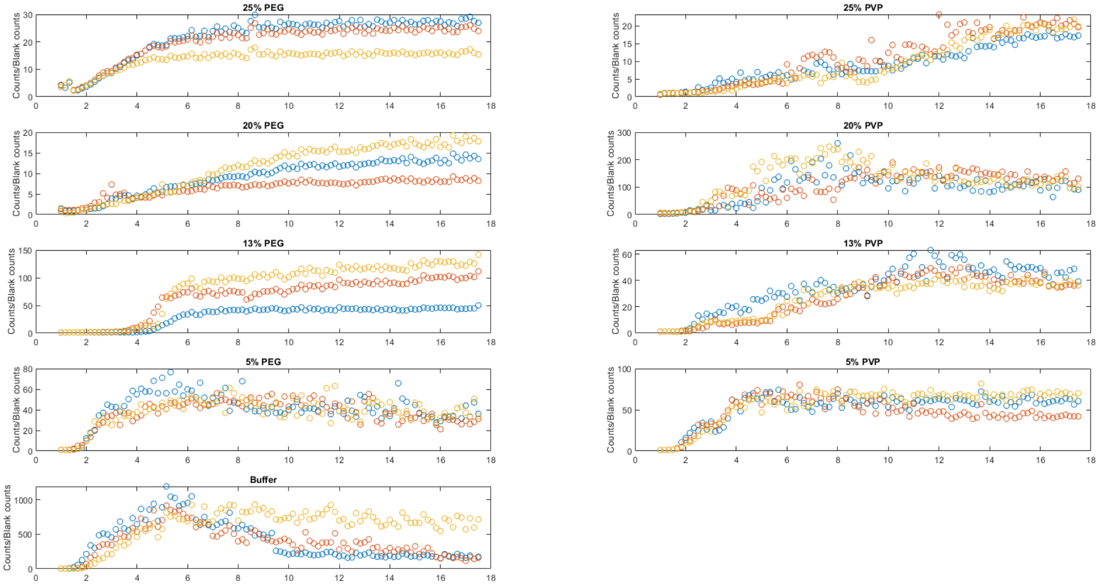


**Figure 6. Thioflavin fluorescent intensity as a function of time for different concentrations of PEG and PVP.**

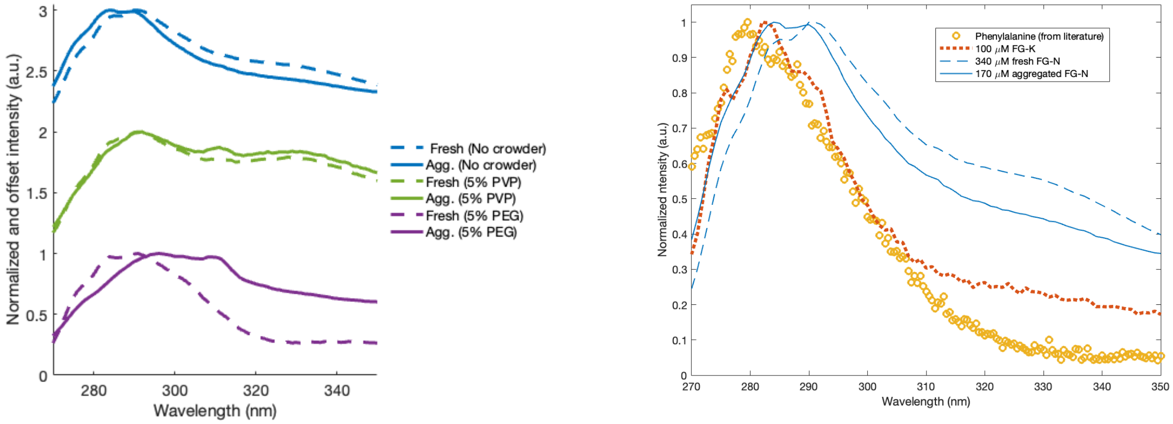


**Figure 7. (left) Phenylalanine fluorescence as a function of aggregation. The same data as presented in Figure 4 of the main text, but with each trace normalized to the same maximum intensity, and offset for ease of comparison. (right) Corresponding data for FG-K is overlayed with the FG-K data (left), as well as the published spectra of phenylalanine in water (https://omlc.org/spectra/PhotochemCAD/html/073.html).**

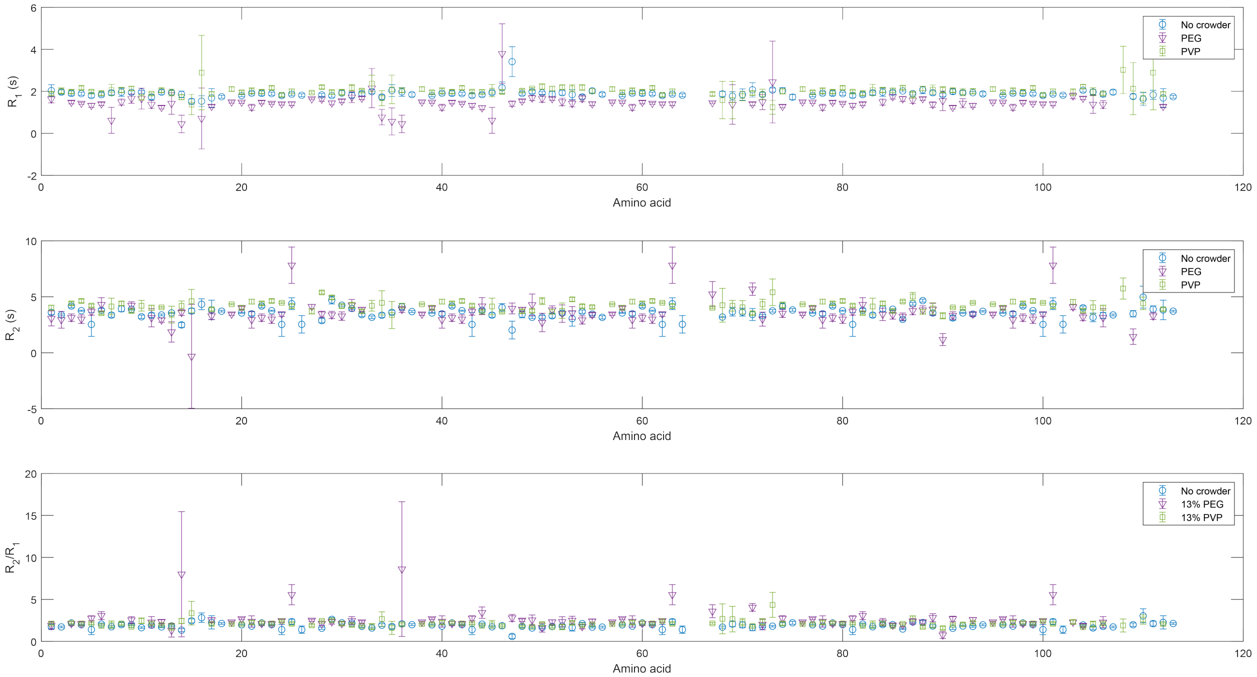


**Figure 8. NMR measured relaxation rates (R1, R2 and R1/R2) in the presence of different crowding agents.**
